# Dissecting Genetic Variability and Multi-Trait Performance in Cucumber

**DOI:** 10.64898/2026.10.30.763243

**Authors:** Mohaimin Islam, Md Salman Shahriar, Mst. Faria Kabir, Md. Anwar Hossain, Md. Omar Kayess

## Abstract

Cucumber improvement requires the simultaneous enhancement of yield, earliness, fruit quality, and postharvest performance, yet many breeding studies rely on single-season or single-trait evaluations. This study therefore assessed 18 diverse cucumber genotypes across two consecutive Rabi seasons (2025 and 2026) using 23 agronomic, phenological, yield, fruit-quality, and postharvest traits, of which 12 non-redundant core traits were retained for the principal genetic and multivariate analyses. Genotypic variability, GCV, PCV, broad-sense heritability, genetic advance, interannual correlation, principal component analysis, hierarchical clustering, and multi-trait selection using MGIDI and MTSI were evaluated. Genotype effects were highly significant for nearly all estimable traits in both years (P<0.001). CU-10 produced the highest yield in both years (6.90 and 7.69 t ha⁻¹), followed by CU-11 (4.38 and 4.89 t ha⁻¹) and CU-09 (4.12 and 4.59 t ha⁻¹). Yield showed high GCV (51.80%), PCV (52.21%), broad-sense heritability (98.46%), and GAM (106.04%). Fruit number plant^−1^ and fruit weight also exhibited high heritability (97.57% and 95.91%) and GAM (80.75% and 59.46%). Interannual repeatability was strongest for fruit number plant^−1^ (r=0.987), yield (r=0.986), and plant mortality (r=0.981), whereas shelf life was less stable (r=0.426, P=0.078). The first two principal components explained 57.29% of total variation, increasing to 70.92% with PC3. MGIDI selected CU-09, CU-11, and check variety Anuska, whereas MTSI selected CU-11, CU-01, and CU-06. Overall, CU-11 was the only genotype selected by both MGIDI and MTSI, indicating a favourable balance of multi-trait performance and relative interannual stability, while the other selected genotypes provided complementary breeding value for hybridization and further validation. These findings provide an integrated framework for improving productivity, stability, and postharvest performance in cucumber breeding programs.

## Introduction

Cucumber (*Cucumis sativus* L., 2n=14) is an economically important cucurbit cultivated across tropical, subtropical, and temperate regions [1,2]. Originating in the Indian subcontinent, it was domesticated from *C. sativus* var. *hardwickii*, generating diverse cultivated germplasm distributed across East Asian, Eurasian, and African gene pools [3,4]. Modern breeding has emphasized gynoecious and parthenocarpic forms for improved productivity and market suitability [3,5]. The adaptability of cucumber permits production under both open-field and protected cultivation systems [6,7].

Cucumber is among the most widely produced vegetable crops globally, with annual production estimated at approximately 91.26–98 million metric tons [3,8]. It ranks among the leading vegetable crops worldwide and has considerable dietary and economic importance [1,7]. Its fruits are widely consumed fresh and processed and provide minerals, vitamins, and antioxidants [1,8]. Growing demand for early, uniform, and high-yielding cultivars has intensified breeding efforts [6]. However, genetic improvement is constrained by the narrow variability of elite cultivars and the poor yield, non-uniformity, unfavorable sex expression, and environmental sensitivity of many unimproved landraces [2,5,6]. Although protected cultivation and optimized crop management can markedly improve productivity [7,9], genetic improvement remains fundamental for achieving sustainable gains in yield and associated traits [10,11,12].

Fruit yield in cucumber is a complex polygenic trait determined by the collective effects of vegetative growth, flowering behavior, reproductive efficiency, and fruit characteristics [1,2]. Considerable genetic variability among cucumber genotypes provides opportunities for effective selection and crop improvement [5,13]. Phenotypic coefficients of variation (PCV) generally exceed genotypic coefficients of variation (GCV), indicating environmental influence on quantitative traits; however, small differences between PCV and GCV suggest substantial genetic control [1,2]. High broad-sense heritability combined with high genetic advance as a percentage of the mean has been reported for flowering, vegetative, and fruit traits, suggesting considerable potential for selection, although broad-sense heritability alone does not distinguish additive from non-additive genetic effects [1,2].

Genetic analyses further indicate that both additive and non-additive gene actions govern yield and associated horticultural traits in cucumber. Additive effects are important for some fruit and quality traits, whereas dominance effects substantially influence fruit yield, fruit number, sex ratio, and flowering behavior, supporting the use of both selection and heterosis breeding strategies [5]. However, breeding decisions based only on individual genetic parameters may be inadequate because yield is influenced by multiple correlated traits. Multivariate approaches, particularly principal component analysis (PCA) and hierarchical clustering, provide complementary information on trait relationships and genetic divergence among genotypes [1,3,13]. PCA can identify major sources of phenotypic variation and influential traits [3,13], whereas cluster analysis can identify genetically divergent genotypes useful for hybridization, heterosis exploitation, and the generation of transgressive segregants [2,3].

Despite these advances, several clear research gaps remain. First, most cucumber variability and genetic-divergence studies have been conducted in a single season or year, restricting assessment of genotype performance across temporal environments and limiting understanding of genotype-by-year interactions [2,7,13]. Consequently, the temporal consistency of genetic variability, heritability, and superior genotype performance remains insufficiently characterized. Second, conventional breeding frequently emphasizes single traits or individual parameters such as GCV, heritability, and genetic advance. Such univariate approaches may produce undesirable correlated responses when important traits exhibit antagonistic relationships or physiological trade-offs [5,12]. Third, although PCA and clustering effectively describe variation and genetic relationships, they do not directly provide an integrated criterion for selecting genotypes that simultaneously approach a desirable multi-trait breeding ideotype. Finally, an integrated two-year assessment combining quantitative genetic parameters, genetic divergence, multivariate analysis, and ideotype-based multi-trait selection remains insufficiently explored in cucumber.

The Multi-Trait Genotype-Ideotype Distance Index (MGIDI) provides a useful framework to address this latter limitation by integrating factor analysis with genotype-to-ideotype distance, reducing multicollinearity among correlated traits and enabling simultaneous selection for several desirable attributes. This approach is particularly relevant in cucumber, where breeding success depends on an appropriate combination of earliness, plant architecture, fruit characteristics, yield components, and overall productivity rather than improvement of a single trait. Integrating MGIDI with conventional quantitative genetic and multivariate analyses may therefore provide a more comprehensive basis for identifying promising genotypes for direct selection, parental choice, hybridization, and population improvement. Accordingly, the present study aimed to evaluate the performance and breeding potential of diverse cucumber genotypes over two consecutive years using an integrated framework of quantitative genetics and multivariate analysis, thereby supporting the selection of genetically diverse and agronomically promising materials for hybrid breeding and future population improvement.

## Materials and Methods

### Experimental Site and Agro-Climatic Conditions

The field experiment was conducted during two consecutive Rabi cropping seasons, in 2025 and 2026 (February–May), at Seed Bangla Agri Genetics, Bogura, Bangladesh (24.8833° N, 89.4500° E; 18 m above sea level). The experimental site is situated within a monsoon-influenced humid subtropical agro-climatic zone. During the Rabi season, weather conditions exhibited a distinct seasonal progression, with increasing temperature, rainfall, and relative humidity from February to May (**Supplementary Table 1**). Mean maximum air temperature increased from 28.1 °C in February to ∼ 33–34 °C during April and May, while mean minimum temperature rose from 15.0 to 24.2 °C. Rainfall remained relatively low during February and March but increased substantially during April and May, reaching 241 mm in May. Relative humidity also increased toward the end of the cropping period, whereas sunshine duration declined during May. These monthly values were derived from weekly normal meteorological observations for standard meteorological weeks 6–22.

### Plant Material and Experimental Design

A panel of 18 diverse cucumber genotypes (14 advanced lines and 4 check varieties) were evaluated (**Supplementary Table 2**). The experiment was laid out in a Randomized Complete Block Design (RCBD) with three replications in each evaluation year. Plants were established in pits spaced 2.0 m × 2.0 m, with a bed-to-bed distance of 0.60 m. Seeds were initially sown in earthen polybags measuring 10.0 cm × 5.5 cm, with two seeds bag^−1^. The nursery medium consisted of well-decomposed cow dung and sandy soil mixed at a 1:1 ratio and solarized before sowing. Raised seed beds approximately 10 cm high were prepared, and 18-day-old healthy seedlings were transplanted into the prepared pits. After establishment, one healthy seedling was maintained pit^−1^ to ensure a uniform plant stand.

### Crop Management and Cultivation Practices

Standard agronomic practices were applied uniformly across all experimental units. The field was prepared by repeated ploughing and laddering, followed by cultivation on raised beds with drainage channels. Well-decomposed farmyard manure (5 t ha⁻¹) and mineral fertilizers comprising urea (80 kg ha⁻¹), TSP (100 kg ha⁻¹), MOP (50 kg ha⁻¹), gypsum (50 kg ha⁻¹), zinc (12 kg ha⁻¹), and boron (10 kg ha⁻¹) were applied during land preparation and crop establishment, with nitrogen supplied in split doses during vegetative growth, flowering, and fruiting. Vines were supported with bamboo stakes. Irrigation was applied according to soil-moisture status, with the first irrigation given about 7–10 days after establishment and subsequent irrigations at approximately 2-week intervals when required. Weeds were controlled manually, including a major weeding operation about 20–25 days after emergence. For pest management, red pumpkin beetle was controlled during the seedling stage, and pheromone traps were installed at one trap per 5 m² during fruiting to manage fruit fly infestation. Cucumber mosaic virus and wilt incidence were monitored periodically.

### Phenotypic Trait Evaluation

Phenotypic evaluation was conducted for 23 agronomic, phenological, yield, fruit-quality, and postharvest traits in both evaluation years. Twelve non-redundant core traits were retained for the principal main-text genetic and multivariate analyses, while results for all 23 traits are provided in the Supplementary Information. The selected traits were those subsequently used for analysis of variance, genetic parameter estimation, correlation analysis, principal component analysis, hierarchical clustering, and multi-trait genotype selection. The 12 core traits comprised plant mortality (PM, %), days to female flowering (DFF), days to first harvest (DFH), fruit length (FL, cm), fruit diameter (FD, cm), number of harvests (NH), average fruit number plant^−1^ (FNPP), average fruit weight (FW, g), fruit yield (YLD, t ha⁻¹), pericarp thickness (PT, cm), shelf life (SL, days), and weight loss after 7 days (WL7, %). Plant mortality was expressed as the percentage of plants that failed to survive during crop establishment. Days to female flowering was recorded as the number of days required for the appearance of the first female flower, whereas days to first harvest represented the time to the first marketable fruit harvest. Fruit length and fruit diameter were recorded in centimeters, and the total number of harvest events was counted throughout the harvesting period. Average fruit number plant^−1^ and average fruit weight were determined for each genotype, while fruit yield was expressed as t ha⁻¹. Fruit quality was further characterized by pericarp thickness. Postharvest performance was assessed using shelf life, expressed in days, and percentage fruit weight loss after 7 days of storage. The same 12 traits were used consistently in the dual-year correlation analysis and multivariate analyses, including PCA and hierarchical clustering, ensuring methodological correspondence between phenotypic characterization and downstream statistical evaluation.

### Genetic Analysis

Genotypic variance (σ²g) and phenotypic variance (σ²p) were estimated according to the standard procedures described by Acquaah [14].

Genotypic variance 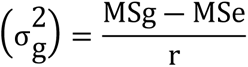

Where MSe is the mean square of the error, MS_g_ is the mean square of genotypes, and “r*”* is the number of replications

Phenotypic Variance (*σ*^2^*p*) = *σ*^2^*g* + *σ*^2^*e*

Here σ^2^p, phenotypic variance; σ^2^g, genotypic variance; σ^2^e, environmental variance as mentioned by Johnson et al. [15].

The genotypic coefficient of variation (GCV) and phenotypic coefficient of variation (PCV) were computed using the equations described by Singh and Chaudhary [16].

Genotypic co ― efficient of variation 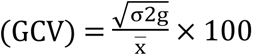

Here,

σ^2^_g_ = Genotypic variance; and

x̄ = Population mean

Phenotypic co ― efficient of variation 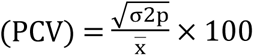

Here,

σ^2^_p_ Phenotypic variance; and

x̄ = Population mean

The magnitudes of GCV and PCV were categorized as low (0–10%), moderate (10– 20%), and high (≥20%), following the classification proposed by Burton and DeVane [17].

Broad-sense heritability (h²b) was estimated as the proportion of genotypic variance (σ²g) to the total phenotypic variance (σ²p), following Falconer and Mackay [18], using the following equation:

Broad-sense heritability was categorized as low (0–30%), moderate (30–60%), and high (≥60%), following the classification proposed by Robinson et al. [19].

Expected genetic advance (GA) and genetic advance as a percentage of the mean (GAM) were estimated following the method of Assefa et al. [20], using a selection intensity (K) corresponding to 5% selection pressure. The GAM values were subsequently classified as low (0–10%), moderate (10–20%), and high (>20%), according to Johnson et al. [15].

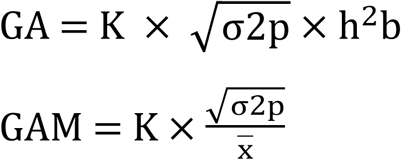

The intensity of the selection is denoted by the constant K. h^2^B stands for heritability, X stands for the mean of the traits, and P is the phenotypic standard deviation when K is 5%.

### Statistical Analysis

Statistical analyses were performed using R software (version 4.2.3). For year-wise analysis, genotype effects were treated as fixed and replication effects as random. Analysis of variance (ANOVA) was conducted under the randomized complete block design to test for significant differences among genotypes. When significant genotype effects were detected, mean separation was performed using Tukey’s honestly significant difference (HSD) test at (P <0.05), consistent with the mean-comparison procedure reported in the genotype tables. For the combined two-year analysis, variance components attributable to genotype, genotype × year interaction, and residual error were estimated and subsequently used for genetic parameter estimation, including the genotypic coefficient of variation (GCV), phenotypic coefficient of variation (PCV), broad-sense heritability, genetic advance, and genetic advance as a percentage of the mean (GAM). Pearson correlation coefficients were calculated among the 12 core traits separately for each year, and interannual consistency was assessed using correlations between genotype means across years. Multivariate relationships among genotypes and traits were further investigated using principal component analysis (PCA) and hierarchical cluster analysis based on standardized trait values, with heatmaps used to visualize multivariate patterns. Multi-trait selection was conducted using the Multi-Trait Genotype–Ideotype Distance Index (MGIDI) and a customized Multi-Trait Stability Index (MTSI) based on 12 core traits across two years. MGIDI used factor analysis and distance from an ideal genotype, while MTSI integrated mean performance and Weighted Average of Absolute Scores (WAAS)-based stability with equal weights (50:50). Genotypes were ranked using both indices, applying a 15% selection intensity. Data processing, statistical computation, and graphical visualization were carried out in R and Microsoft Excel. The R packages metan, ggplot2, FactoMineR, and factoextra were used, as appropriate, for multivariate analysis, clustering, correlation analysis, and graphical presentation.

## Results

### Analysis of variance across years

Analysis of variance revealed pronounced differences among the 18 cucumber genotypes for nearly all evaluated traits in both years (**Tables 1A** and **1B**). In the first year, genotype effects were highly significant (P < 0.001) for plant mortality, days to female flowering, days to first harvest, fruit length, fruit diameter, average fruit number plant⁻¹, average fruit weight, yield, pericarp thickness, and weight loss after 7 days, whereas shelf life also differed significantly among genotypes. A comparable pattern was observed in the second year, with highly significant genotype effects for all estimable traits (P < 0.001). Notably, yield showed strong genotypic differentiation in both years (F = 208.53 and 248.31, respectively), together with substantial differences in major yield components, including average fruit number plant⁻¹ (F = 240.48 and 249.33) and average fruit weight (F = 249.82 and 249.00). Overall, the concurring genotype effects across the two years demonstrate substantial phenotypic variation among the evaluated cucumber genotypes for earliness, fruit attributes, yield components, yield, and postharvest traits.

**Table 1A.**
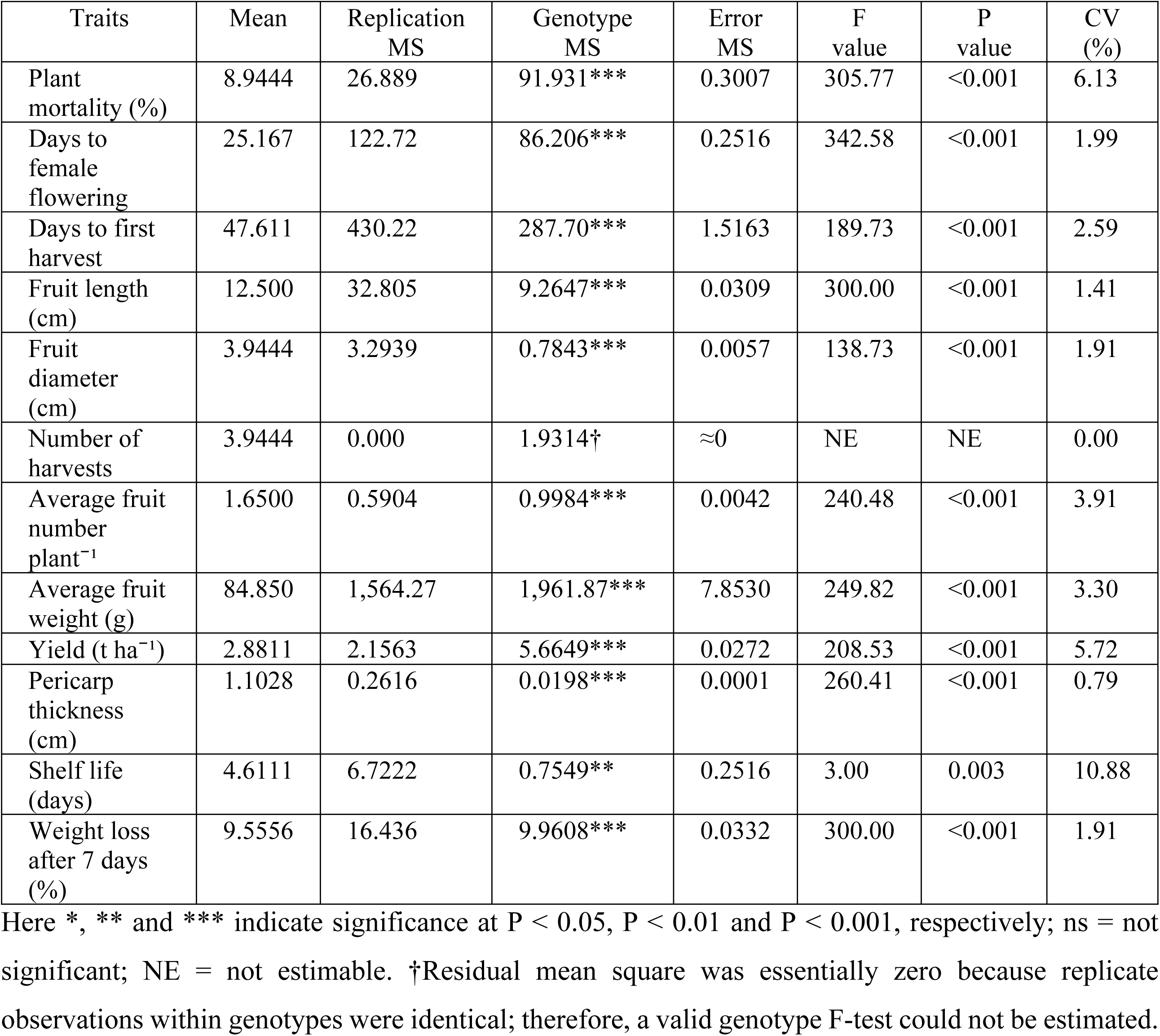
1^st^ year analysis of variance for 12 selected core traits of 18 cucumber genotypes.

**Table 1B.** 2^nd^ year analysis of variance for 12 selected core traits of 18 cucumber genotypes.

| Traits | Mean | Replication MS | Genotype MS | Error MS | F value | P value | CV (%) |
|---|---|---|---|---|---|---|---|
| Plant mortality (%) | 9.2556 | 18.809 | 107.73*** | 0.4524 | 238.12 | <0.001 | 7.27 |
| Days to female flowering | 26.056 | 150.22 | 144.52*** | 0.6928 | 208.60 | <0.001 | 3.19 |
| Days to first harvest | 49.389 | 490.89 | 560.17*** | 2.0654 | 271.22 | <0.001 | 2.91 |
| Fruit length (cm) | 12.233 | 33.347 | 19.514*** | 0.0696 | 280.48 | <0.001 | 2.16 |
| Fruit diameter (cm) | 3.8500 | 3.1250 | 1.9950*** | 0.0074 | 271.32 | <0.001 | 2.23 |
| Number of harvests | 4.0000 | 0.000 | 3.1765† | ≈0 | NE | NE | 0.00 |
| Average fruit number plant<sup>−1</sup> | 1.6589 | 0.5977 | 1.7088*** | 0.0069 | 249.33 | <0.001 | 4.99 |
| Average fruit weight (g) | 82.489 | 1,481.49 | 1,986.06*** | 7.9762 | 249.00 | <0.001 | 3.42 |
| Yield (t ha<sup>−1</sup>) | 2.9022 | 1.8432 | 8.2206*** | 0.0331 | 248.31 | <0.001 | 6.27 |
| Pericarp thickness (cm) | 1.0783 | 0.2640 | 0.1121*** | 0.0004 | 265.51 | <0.001 | 1.91 |
| Shelf life (days) | 4.4444 | 0.8889 | 2.1961*** | 0.1830 | 12.00 | <0.001 | 9.63 |
| Weight loss after 7 days (%) | 9.9167 | 17.602 | 16.123*** | 0.0587 | 274.71 | <0.001 | 2.44 |

Here \*, \*\* and \*\*\* indicate significance at P < 0.05, P < 0.01 and P < 0.001, respectively; ns = not significant; NE = not estimable. Residual mean square was essentially zero because replicate observations within genotypes were identical; therefore, a valid genotype F-test could not be estimated.

### Genotypic mean performance for phenological and fruit traits

Marked genotypic variation was observed in plant mortality, phenological development, fruit morphology, and yield components across both years (**Tables 2A** and **2B**). CU-10 and the check variety Anuska recorded the lowest plant mortality (4.00 and 3.50% in Years 1 and 2, respectively), whereas Green Line consistently exhibited the highest mortality (24.00 and 26.80%). For female flowering, CU-04 and CU-09 were earliest in Year 1 (18 days), while CU-09 remained earliest in Year 2 (16 days), significantly preceding all four check varieties. In contrast, All Rounder exhibited the latest flowering (33 and 37 days), although it was statistically comparable to CU-11 in Year 1 and CU-03 and CU-12 in Year 2. Similarly, CU-05 reached first harvest earliest (38 and 34 days), followed by CU-08 (39 and 35 days) and CU-09 (41 and 36 days), outperforming the checks numerically in both years. Conversely, CU-02, CU-03, and CU-07 required the longest period to first harvest (68 and 76 days), significantly exceeding all checks. These results distinguished CU-05 and CU-09 for early harvest maturity, with CU-09 additionally exhibiting consistently early female flowering.

**Table 2A.**
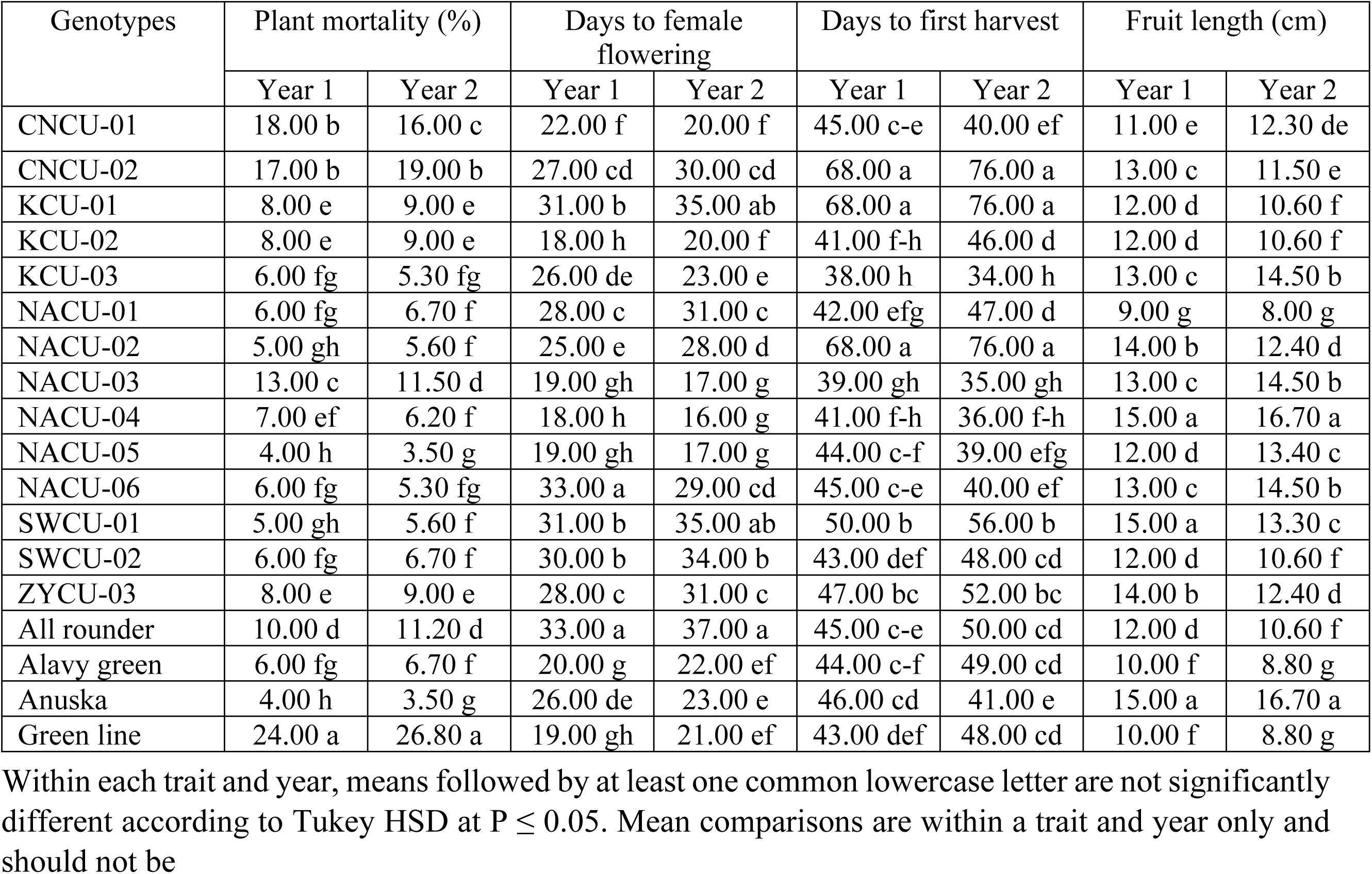
Dual-year genotype means for plant mortality, days to female flowering, days to first harvest and fruit length.

| Genotypes | Plant mortality (%) |  | Days to female flowering |  | Days to first harvest |  | Fruit length (cm) |  |
|---|---|---|---|---|---|---|---|---|
|  | Year 1 | Year 2 | Year 1 | Year 2 | Year 1 | Year 2 | Year 1 | Year 2 |
| CNCU-01 | 18.00 b | 16.00 c | 22.00 f | 20.00 f | 45.00 c-e | 40.00 ef | 11.00 e | 12.30 de |
| CNCU-02 | 17.00 b | 19.00 b | 27.00 cd | 30.00 cd | 68.00 a | 76.00 a | 13.00 c | 11.50 e |
| KCU-01 | 8.00 e | 9.00 e | 31.00 b | 35.00 ab | 68.00 a | 76.00 a | 12.00 d | 10.60 f |
| KCU-02 | 8.00 e | 9.00 e | 18.00 h | 20.00 f | 41.00 f-h | 46.00 d | 12.00 d | 10.60 f |
| KCU-03 | 6.00 fg | 5.30 fg | 26.00 de | 23.00 e | 38.00 h | 34.00 h | 13.00 c | 14.50 b |
| NACU-01 | 6.00 fg | 6.70 f | 28.00 c | 31.00 c | 42.00 efg | 47.00 d | 9.00 g | 8.00 g |
| NACU-02 | 5.00 gh | 5.60 f | 25.00 e | 28.00 d | 68.00 a | 76.00 a | 14.00 b | 12.40 d |
| NACU-03 | 13.00 c | 11.50 d | 19.00 gh | 17.00 g | 39.00 gh | 35.00 gh | 13.00 c | 14.50 b |
| NACU-04 | 7.00 ef | 6.20 f | 18.00 h | 16.00 g | 41.00 f-h | 36.00 f-h | 15.00 a | 16.70 a |
| NACU-05 | 4.00 h | 3.50 g | 19.00 gh | 17.00 g | 44.00 c-f | 39.00 efg | 12.00 d | 13.40 c |
| NACU-06 | 6.00 fg | 5.30 fg | 33.00 a | 29.00 cd | 45.00 c-e | 40.00 ef | 13.00 c | 14.50 b |
| SWCU-01 | 5.00 gh | 5.60 f | 31.00 b | 35.00 ab | 50.00 b | 56.00 b | 15.00 a | 13.30 c |
| SWCU-02 | 6.00 fg | 6.70 f | 30.00 b | 34.00 b | 43.00 def | 48.00 cd | 12.00 d | 10.60 f |
| ZYCU-03 | 8.00 e | 9.00 e | 28.00 c | 31.00 c | 47.00 bc | 52.00 bc | 14.00 b | 12.40 d |
| All rounder | 10.00 d | 11.20 d | 33.00 a | 37.00 a | 45.00 c-e | 50.00 cd | 12.00 d | 10.60 f |
| Alavy green | 6.00 fg | 6.70 f | 20.00 g | 22.00 ef | 44.00 c-f | 49.00 cd | 10.00 f | 8.80 g |
| Anuska | 4.00 h | 3.50 g | 26.00 de | 23.00 e | 46.00 cd | 41.00 e | 15.00 a | 16.70 a |
| Green line | 24.00 a | 26.80 a | 19.00 gh | 21.00 ef | 43.00 def | 48.00 cd | 10.00 f | 8.80 g |

Within each trait and year, means followed by at least one common lowercase letter are not significantly different according to Tukey HSD at P ≤ 0.05. Mean comparisons are within a trait and year only and should not be

**Table 2B.** Dual-year genotype means for fruit diameter, number of harvests, average fruit number per plant and average fruit weight.

| Genotypes | Fruit diameter (cm) |  | Number of harvests |  | Average fruit number plant<sup>−1</sup> |  | Average fruit weight (g) |  |
|---|---|---|---|---|---|---|---|---|
|  | Year 1 | Year 2 | Year 1 | Year 2 | Year 1 | Year 2 | Year 1 | Year 2 |
| CNCU-01 | 5.00 a | 5.60 a | 4.00 | 4.00 | 2.00 d | 2.23 cd | 56.00 h | 62.40 h |
| CNCU-02 | 4.00 c | 3.50 e | 3.00 | 3.00 | 0.92 i | 0.81 i | 135.90 a | 120.30 b |
| KCU-01 | 4.00 c | 3.50 e | 3.00 | 3.00 | 0.85 i | 0.75 i | 107.80 cd | 95.40 e |
| KCU-02 | 4.00 c | 3.50 e | 5.00 | 4.00 | 1.91 d | 1.69 e | 64.80 g | 57.40 hi |
| KCU-03 | 3.50 d | 3.90 d | 4.00 | 4.00 | 2.21 c | 2.47 bc | 65.83 fg | 73.40 g |
| NACU-01 | 3.50 d | 3.10 f | 4.00 | 4.00 | 1.00 hi | 0.88 hi | 104.23 cd | 92.30 e |
| NACU-02 | 4.00 c | 3.50 e | 4.00 | 4.00 | 1.26 fg | 1.11 gh | 111.47 c | 98.60 de |
| NACU-03 | 4.00 c | 4.50 c | 4.00 | 4.00 | 2.33 c | 2.60 b | 66.67 fg | 74.30 fg |
| NACU-04 | 4.50 b | 5.00 b | 5.00 | 6.00 | 2.61 b | 2.91 a | 74.30 f | 82.90 f |
| NACU-05 | 3.50 d | 3.90 d | 6.00 | 7.00 | 2.81 a | 3.13 a | 94.03 e | 104.80 cd |
| NACU-06 | 5.00 a | 5.60 a | 4.00 | 4.00 | 1.83 d | 2.04 d | 122.17 b | 136.20 a |
| SWCU-01 | 4.00 c | 3.50 e | 4.00 | 4.00 | 1.37 ef | 1.21 fg | 111.00 c | 98.20 de |
| SWCU-02 | 4.00 c | 3.50 e | 4.00 | 4.00 | 1.35 f | 1.19 fg | 65.00 g | 57.50 hi |
| ZYCU-03 | 4.00 c | 3.50 e | 3.00 | 3.00 | 1.33 f | 1.18 fg | 62.50 gh | 55.30 hi |
| All rounder | 4.00 c | 3.50 e | 3.00 | 3.00 | 1.40 ef | 1.24 fg | 61.40 gh | 54.40 hi |
| Alavy green | 3.50 d | 3.10 f | 4.00 | 4.00 | 1.13 gh | 1.00 ghi | 65.10 g | 57.60 hi |
| Anuska | 3.50 d | 3.90 d | 4.00 | 4.00 | 1.83 d | 2.04 d | 99.70 de | 111.20 c |
| Green line | 3.00 e | 2.70 g | 3.00 | 3.00 | 1.56 e | 1.38 f | 59.40 gh | 52.60 i |

Within each trait and year, means followed by at least one common lowercase letter are not significantly different according to Tukey HSD at P ≤ 0.05. Mean comparisons are within a trait and year only and should not be interpreted as direct tests between years.

Fruit morphology revealed contrasting performance among genotypes and check varieties. CU-09 and Anuska produced the longest fruits (15.00 and 16.70 cm), significantly surpassing All Rounder, Alavy Green, and Green Line in both years, whereas CU-06 recorded the shortest fruits among the experimental genotypes (9.00 and 8.00 cm). In contrast, CU-01 and CU-11 consistently produced the greatest fruit diameter (5.00 and 5.60 cm), significantly above all four checks, while Green Line exhibited the smallest diameter (3.00 and 2.70 cm). Thus, CU-09 matched Anuska for fruit length, whereas CU-01 and CU-11 surpassed the checks in fruit diameter.

For yield components, CU-10 recorded the highest numerical number of harvests (6 and 7), followed by CU-09 (5 and 6), compared with 3–4 harvests among the checks. CU-10 also produced the highest average fruit number per plant (2.81 and 3.13), followed by CU-09 (2.61 and 2.91), both exceeding the best-performing check, Anuska (1.83 and 2.04), although CU-09 and CU-10 were statistically comparable in Year 2. CU-08 (2.33 and 2.60) and CU-05 (2.21 and 2.47) also surpassed all four checks numerically, whereas CU-02 and CU-03 consistently produced fewer than one fruit per plant. Conversely, CU-02 recorded the highest individual fruit weight in Year 1 (135.90 g), while CU-11 ranked highest in Year 2 (136.20 g), both significantly exceeding the check varieties in their respective years. CU-11 also maintained relatively high fruit weight in Year 1 (122.17 g), whereas Green Line recorded the lowest numerical values (59.40 and 52.60 g). Overall, CU-10 combined low mortality with high fruit number and harvest frequency, CU-09 combined early flowering and harvest maturity with long fruits and high fruit number, and CU-11 was distinguished by greater fruit diameter and individual fruit weight relative to the check varieties.

### Yield and postharvest performance

Marked genotypic differences were observed for yield, pericarp thickness, and postharvest weight loss, whereas shelf-life responses were less differentiated, particularly in the first year (**Table 3**). CU-10 was the highest-yielding genotype in both years, producing 6.90 and 7.69 t ha⁻¹, respectively. The next highest yields were recorded for CU-11 (4.38 and 4.89 t ha⁻¹) and CU-09 (4.12 and 4.59 t ha⁻¹), while CU-03 consistently produced the lowest yield (1.14 and 1.01 t ha⁻¹). Check variety Anuska also maintained relatively high productivity across years (3.58 and 3.99 t ha⁻¹). Pericarp thickness also varied substantially among genotypes. CU-11 had the thickest pericarp in Year 1 (1.25 cm) and remained in the highest statistical group in Year 2 (1.39 cm), together with CU-01, CU-05, CU-08, and CU-09. In contrast, CU-03, CU-06, CU-13, and All-rounder were among the thinnest-pericarp genotypes in both years. Shelf-life response differed between years. Although the overall ANOVA indicated significant genotypic variation in Year 1, Tukey HSD did not resolve pairwise differences among individual genotype means. In Year 2, clearer genotypic separation was observed, with CU-01, CU-05, CU-08, and check variety Anuska showing superior shelf-life performance relative to the remaining genotypes. More pronounced differences were evident for weight loss after 7 days, where lower values indicate better moisture retention. CU-06 recorded the lowest weight loss across the two years (6.00 and 6.20%), followed closely by CU-11 (7.00 and 6.80%); CU-01 and check variety Anuska also maintained relatively low losses. By contrast, CU-13 exhibited the greatest weight loss (13.00 and 14.80%), with check variety Alavy green also showing consistently high losses (12.00 and 13.60%). CU-10 combined exceptionally high yield with high fruit number but only moderate postharvest performance, whereas CU-11 combined high yield, heavy fruits, a thick pericarp, and relatively low postharvest weight loss.

**Table 3.** Dual-year genotype means for yield, pericarp thickness, shelf life and weight loss after 7 days.

| Genotypes | Yield (t ha<sup>−1</sup>) |  | Pericarp thickness (cm) |  | Shelf life (days) |  | Weight loss after 7 days (%) |  |
|---|---|---|---|---|---|---|---|---|
|  | Year 1 | Year 2 | Year 1 | Year 2 | Year 1 | Year 2 | Year 1 | Year 2 |
| CNCU-01 | 2.28 gh | 2.54 fg | 1.20 b | 1.34 a | 5.00 a | 6.00 a | 8.00 f | 7.20 i |
| CNCU-02 | 2.88 ef | 2.55 fg | 1.10 c | 0.97 d | 5.00 a | 4.00 b | 10.00 d | 11.40 d |
| KCU-01 | 1.14 k | 1.01 k | 1.00 d | 0.88 e | 5.00 a | 4.00 b | 9.00 e | 10.20 ef |
| KCU-02 | 2.43 fg | 2.15 gh | 1.10 c | 0.97 d | 5.00 a | 4.00 b | 11.00 c | 12.50 c |
| KCU-03 | 3.30 cde | 3.68 cd | 1.20 b | 1.34 a | 5.00 a | 6.00 a | 10.00 d | 9.00 gh |
| NACU-01 | 2.04 ghi | 1.80 hi | 1.00 d | 0.88 e | 4.00 a | 4.00 b | 6.00 h | 6.20 j |
| NACU-02 | 3.45 cd | 3.05 ef | 1.10 c | 0.97 d | 5.00 a | 4.00 b | 8.00 f | 9.10 gh |
| NACU-03 | 3.04 de | 3.39 de | 1.20 b | 1.34 a | 5.00 a | 6.00 a | 12.00 b | 10.80 de |
| NACU-04 | 4.12 b | 4.59 b | 1.20 b | 1.34 a | 4.00 a | 4.00 b | 9.00 e | 8.70 h |
| NACU-05 | 6.90 a | 7.69 a | 1.00 d | 1.12 c | 4.00 a | 4.00 b | 10.00 d | 9.70 fg |
| NACU-06 | 4.38 b | 4.89 b | 1.25 a | 1.39 a | 4.00 a | 4.00 b | 7.00 g | 6.80 ij |
| SWCU-01 | 1.99 ghi | 1.76 hij | 1.10 c | 0.97 d | 4.00 a | 4.00 b | 9.00 e | 9.30 gh |
| SWCU-02 | 2.01 ghi | 1.78 hij | 1.00 d | 0.88 e | 5.00 a | 4.00 b | 13.00 a | 14.80 a |
| ZYCU-03 | 1.63 ijk | 1.44 ijk | 1.10 c | 0.97 d | 4.00 a | 4.00 b | 11.00 c | 11.40 d |
| All rounder | 1.40 jk | 1.24 jk | 1.00 d | 0.88 e | 4.00 a | 4.00 b | 10.00 d | 10.40 ef |
| Alavy green | 1.80 hij | 1.60 hij | 1.10 c | 0.97 d | 5.00 a | 4.00 b | 12.00 b | 13.60 b |
| Anuska | 3.58 c | 3.99 c | 1.10 c | 1.23 b | 5.00 a | 6.00 a | 8.00 f | 7.20 i |
| Green line | 3.49 cd | 3.09 ef | 1.10 c | 0.97 d | 5.00 a | 4.00 b | 9.00 e | 10.20 ef |

Within each trait and year, means followed by at least one common lowercase letter are not significantly different according to Tukey HSD at P ≤ 0.05. Mean comparisons are within a trait and year only and should not be interpreted as direct tests between years.

### Genetic analysis of the selected traits of cucumber genotypes evaluated over two years

Considerable genetic variability was observed for most traits, with PCV consistently exceeding GCV, although the differences were generally small (**Table 4**). The closest GCV–PCV estimates occurred for plant mortality (62.68 vs. 63.04%), yield (51.80 vs. 52.21%), fruit number plant⁻¹ (39.62 vs. 40.12%), fruit weight (29.43 vs. 30.05%), and weight loss after 7 days (20.21 vs. 20.82%). Together with high broad-sense heritability (94.22–98.88%), these results indicate that variation in these traits was largely genetically determined, with relatively limited environmental influence.

**Table 4.**
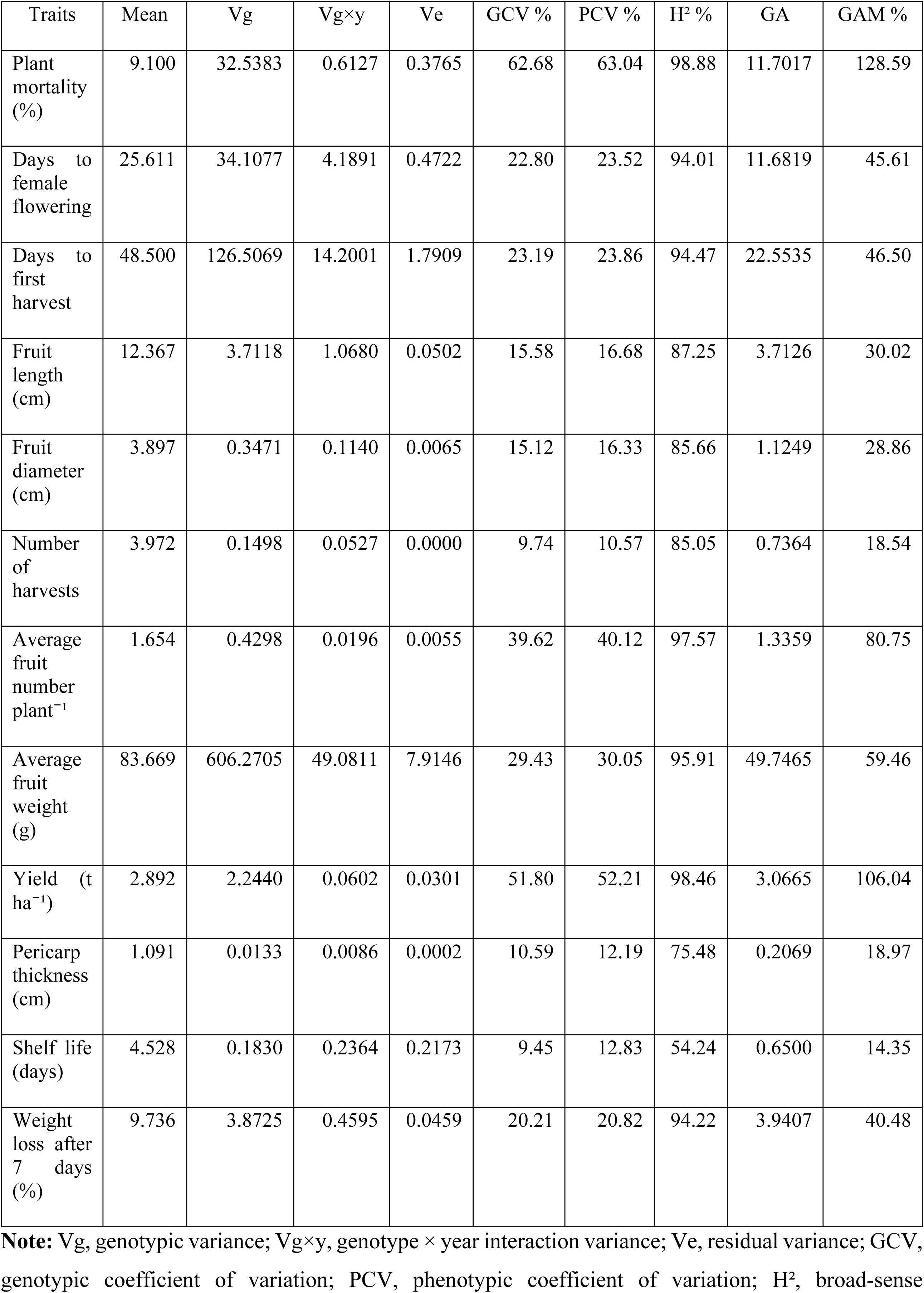
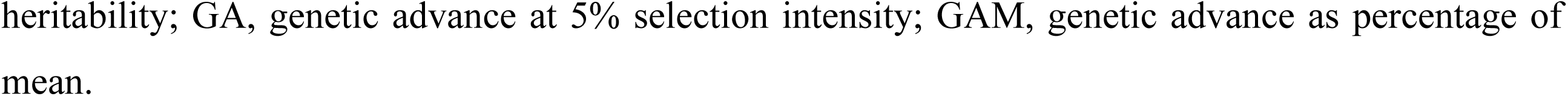
Genetic analysis for 12 selected traits of cucumber genotypes evaluated over two years.

| Traits | Mean | Vg | Vg×y | Ve | GCV % | PCV % | H<sup>2</sup> % | GA | GAM % |
|---|---|---|---|---|---|---|---|---|---|
| Plant mortality (%) | 9.100 | 32.5383 | 0.6127 | 0.3765 | 62.68 | 63.04 | 98.88 | 11.7017 | 128.59 |
| Days to female flowering | 25.611 | 34.1077 | 4.1891 | 0.4722 | 22.80 | 23.52 | 94.01 | 11.6819 | 45.61 |
| Days to first harvest | 48.500 | 126.5069 | 14.2001 | 1.7909 | 23.19 | 23.86 | 94.47 | 22.5535 | 46.50 |
| Fruit length (cm) | 12.367 | 3.7118 | 1.0680 | 0.0502 | 15.58 | 16.68 | 87.25 | 3.7126 | 30.02 |
| Fruit diameter (cm) | 3.897 | 0.3471 | 0.1140 | 0.0065 | 15.12 | 16.33 | 85.66 | 1.1249 | 28.86 |
| Number of harvests | 3.972 | 0.1498 | 0.0527 | 0.0000 | 9.74 | 10.57 | 85.05 | 0.7364 | 18.54 |
| Average fruit number plant<sup>−1</sup> | 1.654 | 0.4298 | 0.0196 | 0.0055 | 39.62 | 40.12 | 97.57 | 1.3359 | 80.75 |
| Average fruit weight (g) | 83.669 | 606.2705 | 49.0811 | 7.9146 | 29.43 | 30.05 | 95.91 | 49.7465 | 59.46 |
| Yield (t ha<sup>−1</sup>) | 2.892 | 2.2440 | 0.0602 | 0.0301 | 51.80 | 52.21 | 98.46 | 3.0665 | 106.04 |
| Pericarp thickness (cm) | 1.091 | 0.0133 | 0.0086 | 0.0002 | 10.59 | 12.19 | 75.48 | 0.2069 | 18.97 |
| Shelf life (days) | 4.528 | 0.1830 | 0.2364 | 0.2173 | 9.45 | 12.83 | 54.24 | 0.6500 | 14.35 |
| Weight loss after 7 days (%) | 9.736 | 3.8725 | 0.4595 | 0.0459 | 20.21 | 20.82 | 94.22 | 3.9407 | 40.48 |

**Note:** Vg, genotypic variance; Vg×y, genotype × year interaction variance; Ve, residual variance; GCV, genotypic coefficient of variation; PCV, phenotypic coefficient of variation; H<sup>2</sup>, broad-sense heritability; GA, genetic advance at 5% selection intensity; GAM, genetic advance as percentage of mean.

The greatest scope for selection was observed for yield, fruit number plant⁻¹, and fruit weight, which combined high genetic variability, heritability, and genetic advance as percent of mean (GAM). Yield recorded 98.46% heritability and 106.04% GAM, fruit number plant⁻¹ 97.57% heritability and 80.75% GAM, and fruit weight 95.91% heritability and 59.46% GAM, indicating strong potential for improvement through phenotypic selection. Days to female flowering and days to first harvest also showed high heritability (94.01–94.47%) and GAM (45.61–46.50%), suggesting good prospects for selection for earliness. Fruit length and diameter showed high heritability and high GAM, although their expected genetic gain was lower than that observed for the major yield components. Plant mortality showed the highest variability (GCV = 62.68%; PCV = 63.04%), heritability (98.88%), and GAM (128.59%). Because lower mortality is desirable, this variability provides substantial opportunity for selecting genotypes with reduced plant loss. Weight loss after 7 days also exhibited high heritability (94.22%) and appreciable GAM (40.48%), indicating potential for improving postharvest moisture retention.

In contrast, shelf life showed the weakest genetic control, with low GCV (9.45%), the lowest heritability (54.24%), and GAM of 14.35%. Its genotype × year variance (0.2364) exceeded the genotypic variance (0.1830), indicating considerable year-dependent expression. Pericarp thickness also showed comparatively greater environmental sensitivity. These traits therefore require evaluation across environments or seasons before reliable selection. Overall, yield, fruit number plant⁻¹, fruit weight, earliness, and postharvest weight loss emerged as the most promising selection targets. Their high heritability and genetic advance, together with relatively small genotype × year effects, indicate comparatively stable genetic expression, whereas shelf life, pericarp thickness, fruit length, and fruit diameter showed greater environmental modulation and warrant multi-environment validation.

### Interannual consistency of trait expression

Genotype performance was highly consistent across the two years for most traits (**Table 5**). Eleven of the twelve traits showed highly significant positive Year 1–Year 2 correlations (P < 0.001), ranging from r = 0.828 to 0.987. The strongest repeatability was observed for fruit number plant⁻¹ (r = 0.987), yield (r = 0.986), and plant mortality (r = 0.981), indicating highly stable genotype rankings across years. Days to first harvest (r = 0.945), number of harvests (r = 0.926), fruit weight (r = 0.921), days to female flowering (r = 0.917), and weight loss after 7 days (r = 0.917) were also strongly repeatable. Fruit length (r = 0.828), fruit diameter (r = 0.832), and pericarp thickness (r = 0.848) showed significant but comparatively lower interannual consistency, consistent with their larger genotype × year variance components. Their genotype rankings were broadly maintained, although trait expression was more responsive to annual environmental conditions. Shelf life was the only clear exception, with a moderate, non-significant correlation between years (r = 0.426, P = 0.078). This agrees with its lower heritability and relatively large genotype × year variance, indicating greater sensitivity to year-specific conditions. Overall, genotype rankings were highly repeatable for yield, fruit number plant⁻¹, plant mortality, phenological traits, fruit weight, and postharvest weight loss, supporting their reliability for selection across years. In contrast, shelf life requires evaluation across additional seasons or environments for dependable genotype discrimination.

**Table 5.** Interannual consistency of 12 selected cucumber traits based on correlations between genotype means in Year 1 and Year 2.

| Traits | Pearson r | P-value | Significance |
|---|---|---|---|
| Plant mortality (%) | 0.981 | <0.001 | *** |
| Days to female flowering | 0.917 | <0.001 | *** |
| Days to first harvest | 0.945 | <0.001 | *** |
| Fruit length (cm) | 0.828 | <0.001 | *** |
| Fruit diameter (cm) | 0.832 | <0.001 | *** |
| Number of harvests | 0.926 | <0.001 | *** |
| Average fruit number plant<sup>−1</sup> | 0.987 | <0.001 | *** |
| Average fruit weight (g) | 0.921 | <0.001 | *** |
| Yield (t ha<sup>−1</sup>) | 0.986 | <0.001 | *** |
| Pericarp thickness (cm) | 0.848 | <0.001 | *** |
| Shelf life (days) | 0.426 | 0.078 | ns |
| Weight loss after 7 days (%) | 0.917 | <0.001 | *** |

### Trait association patterns across years

Pearson correlation analysis revealed a coherent but not entirely identical correlation structure across the two years (**Figure 1**). In both seasons, yield (YLD) was associated more strongly with fruit production dynamics than with fruit size alone. Most notably, YLD showed consistently strong positive correlations with average fruit number plant^−1^ (FNPP) and number of harvests (NH), with coefficients of 0.72 and 0.68 in Year 1 and 0.81 and 0.79 in Year 2, respectively. By contrast, the association between YLD and fruit weight (FW) was comparatively weak in Year 1 (r = 0.21) and only moderate in Year 2 (r = 0.47).

**Fig 1.**
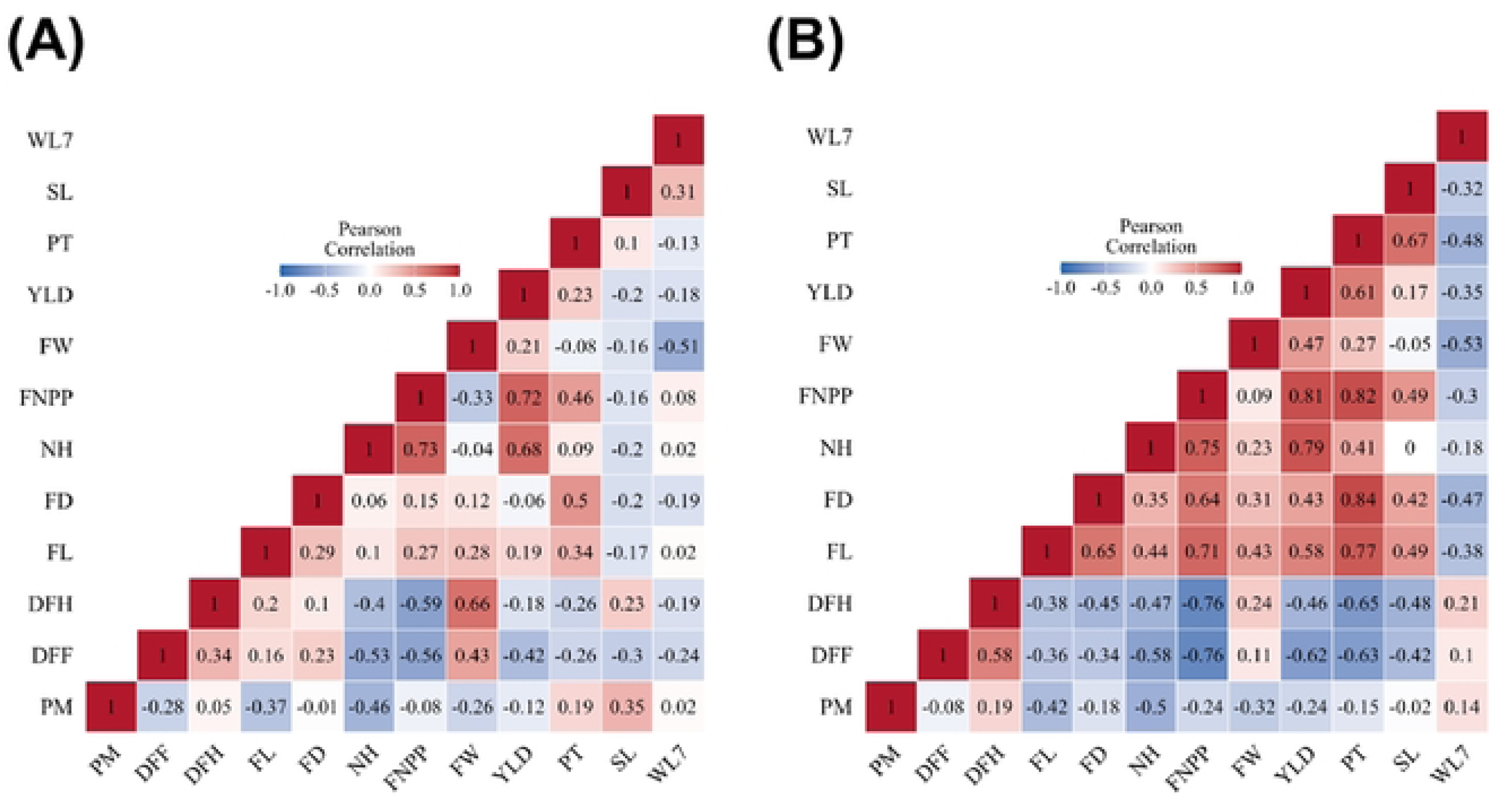
Pearson’s correlation analysis illustrating the relationships among 12 agronomic, yield, and postharvest traits of 18 cucumber genotypes evaluated over two consecutive years. (A) Correlation heatmap showing the relationships among the studied traits in Year 1. (B) Correlation heatmap illustrating the corresponding trait associations in Year 2. The numerical values within each cell represent Pearson’s correlation coefficients (r), indicating the direction and strength of pairwise trait associations. The color gradient ranges from −1 to +1, with red indicating positive correlations, blue indicating negative correlations, and white representing weak or near-zero correlations. Differences in correlation patterns between the two years highlight year-to-year variation in trait associations. **Note:** PM: Plant mortality; DFF: Days to female flowering; DFH: Days to first harvest; FL: Fruit length; FD: Fruit diameter; NH: Number of harvests; FNPP: Average fruit number per plant; FW: Average fruit weight; YLD: Yield; PT: Pericarp thickness; SL: Shelf life; WL7: Weight loss after 7 days.

Phenological traits showed similarly informative relationships with productivity. In both years, days to female flowering (DFF) and days to first harvest (DFH) were negatively associated with FNPP, NH, and YLD, and these negative relationships became more pronounced in Year 2. For example, DFF correlated with FNPP and YLD at −0.56 and −0.42 in Year 1, but at −0.76 and −0.62 in Year 2, respectively. DFH followed the same tendency, especially with FNPP (−0.59 in Year 1 and −0.76 in Year 2). Biologically, this indicates that earlier genotypes tended to produce more fruits and achieve greater yield, whereas delayed flowering and harvesting were generally associated with reduced production efficiency. At the same time, DFF and DFH were positively related to FW, particularly in Year 1 (0.43 and 0.66, respectively), suggesting a partial trade-off in which later genotypes tended to bear fewer but heavier fruits.

Fruit morphology and structural traits became more tightly integrated with yield in Year 2 than in Year 1. In the first year, fruit length (FL) and fruit diameter (FD) showed only weak associations with YLD (0.19 and −0.06, respectively), whereas in the second year both traits were positively related to YLD (0.58 and 0.43). Likewise, pericarp thickness (PT) was only weakly associated with YLD in Year 1 (r = 0.23), but showed a much stronger positive relationship in Year 2 (r = 0.61). PT was also strongly correlated with FL and FD, particularly in Year 2 (r = 0.77 and 0.84), indicating that genotypes with larger fruits also tended to have a thicker pericarp. Moreover, FNPP was positively associated with PT in both years (0.46 and 0.82), reinforcing the idea that some high-performing genotypes combined prolific fruiting with desirable structural fruit attributes.

Postharvest relationships were comparatively less stable across years, which is consistent with the weaker interannual repeatability previously observed for shelf life (SL). Weight loss after 7 days (WL7) was consistently negatively associated with FW in both years (−0.51 and −0.53), implying that heavier fruits tended to lose proportionally less weight during storage. However, the behavior of SL differed between years: in Year 1 it showed only weak relationships with most traits and was even slightly positively associated with WL7 (r = 0.31), whereas in Year 2 it was positively associated with PT (r = 0.67), FL (r = 0.49), FD (r = 0.42), and FNPP (r = 0.49), while being negatively associated with WL7 (r = −0.32). These shifts suggest that postharvest performance was more environmentally contingent than core yield traits, and that structural characteristics such as pericarp thickness may contribute more clearly to storability under some seasonal conditions than others. Another biologically meaningful pattern was the generally negative association of plant mortality (PM) with most productive traits, especially NH, FNPP, FW, and YLD. These relationships were mostly modest in Year 1 but became clearer in Year 2, again indicating that better plant establishment and survival tended to support stronger reproductive performance. Taken together, the correlation network points to a practical breeding message: yield improvement in this germplasm is likely to be most effectively achieved through simultaneous selection for earliness, higher fruit number, and repeated harvest capacity, while postharvest quality traits especially shelf life should be evaluated more cautiously because their trait associations appear less stable across years.

### Principal component analysis based on pooled performance across two years

Principal component analysis based on pooled adjusted genotype means revealed substantial multivariate differentiation among the 18 cucumber genotypes for the 12 core traits (**Figure 2**). The first two principal components explained 57.29% of the total standardized variation, with PC1 and PC2 accounting for 38.23% and 19.06%, respectively. In the two-dimensional genotype–trait biplot (**Figure 2A**), yield (YLD), average fruit number plant^−1^ (FNPP), number of harvests (NH), fruit length (FL), fruit diameter (FD), and pericarp thickness (PT) were oriented broadly toward the same region of the ordination, indicating coordinated contributions to the productive phenotype. CU-09 and CU-10 occupied the side of the biplot associated with YLD, FNPP and NH, whereas CU-13, CU-02 and CU-06 were positioned more closely toward the phenological vectors DFF and DFH.

**Fig 2.**
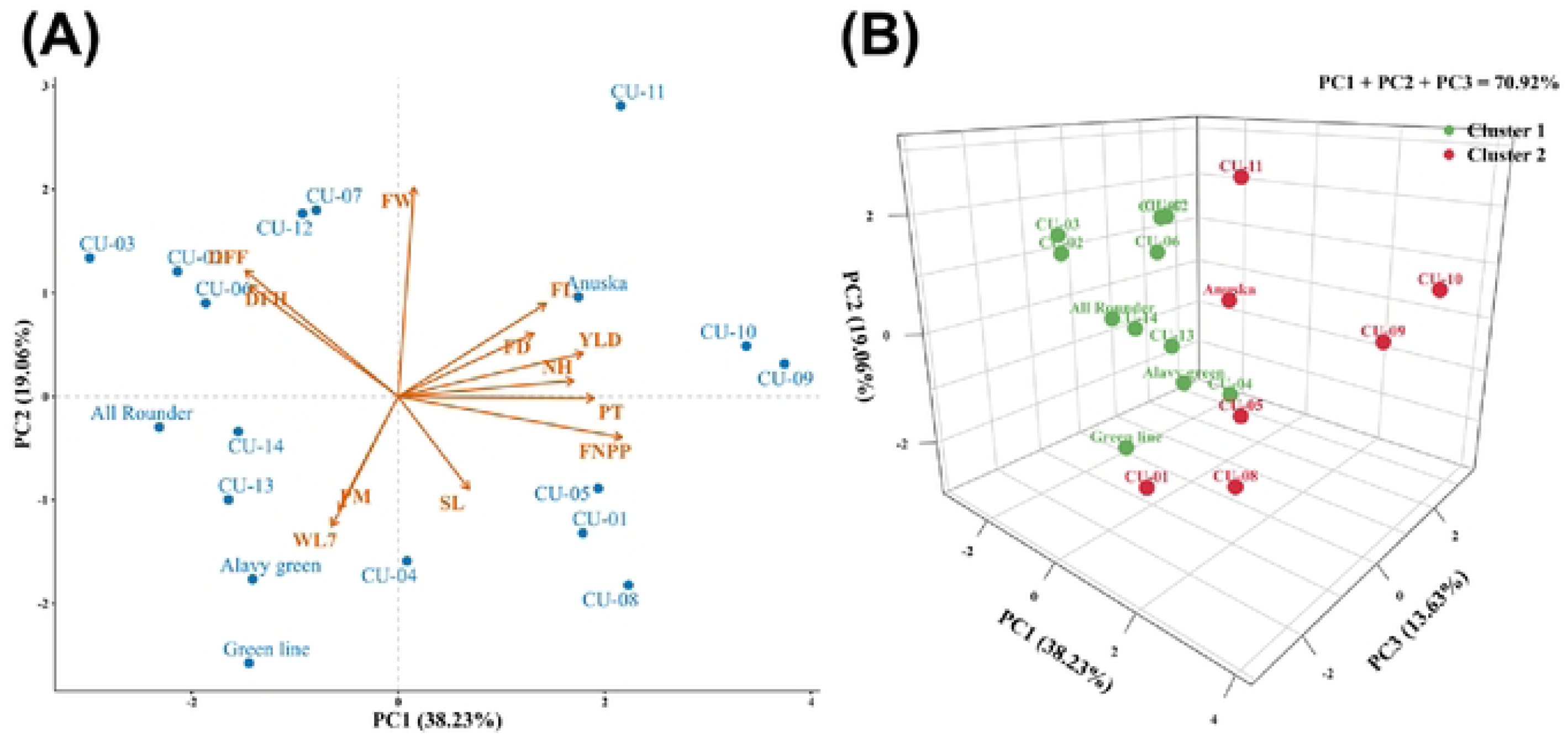
Principal component analysis (PCA) illustrating the multivariate relationships and clustering patterns of 18 cucumber genotypes based on 12 agronomic, yield, and postharvest traits. (A) Two-dimensional PCA biplot showing the distribution of cucumber genotypes and their associations with the studied traits along the first two principal components, PC1 (38.23%) and PC2 (19.06%), which collectively explained 57.29% of the total variation. Blue points represent individual genotypes, while orange arrows indicate trait vectors, with their directions and lengths reflecting the associations of traits with the principal components. (B) Three-dimensional PCA score plot illustrating the distribution of cucumber genotypes along PC1 (38.23%), PC2 (19.06%), and PC3 (13.63%), collectively accounting for 70.92% of the total variation. Green and red points represent Clusters 1 and 2, respectively, as identified independently through hierarchical cluster analysis. The spatial distribution of genotypes highlights their multivariate similarities and differences based on the evaluated traits.

Inclusion of PC3 increased the cumulative explained variation to 70.92%, with PC3 contributing 13.63% (**Figure 2B**). The three-dimensional score plot provided additional separation among genotypes and showed that the two groups identified independently by hierarchical cluster analysis occupied broadly distinct regions of PCA space. Several high-performing genotypes, including CU-09, CU-10 and CU-11, belonged to Cluster 2, whereas Cluster 1 contained genotypes representing contrasting combinations of phenology, productivity and postharvest characteristics. Thus, the 2D and 3D PCA visualizations jointly demonstrated that multivariate differentiation among genotypes was primarily associated with coordinated variation in earliness, fruiting intensity, fruit morphology, yield and postharvest traits.

### Hierarchical clustering of cucumber genotypes based on pooled trait profiles

Hierarchical clustering of standardized pooled adjusted means separated the cucumber genotypes into two major clusters (**Figure 3**), reflecting differences in combined agronomic, yield, fruit morphology, phenology, and postharvest traits. The smaller Cluster 2 comprised CU-01, check variety Anuska, CU-05, CU-08, CU-11, CU-09, and CU-10 and generally showed superior productivity, with above-average number of harvests, fruit number plant⁻¹, and yield. CU-09 and CU-10 were particularly associated with prolific fruiting and high yield, whereas CU-11 combined strong productivity with higher fruit weight and thicker pericarp. CU-01 and check variety Anuska showed relatively favorable postharvest profiles, indicating that Cluster 2 contained several complementary high-performing trait combinations rather than a single uniform ideotype. The larger Cluster 1 contained CU-06, CU-07, CU-12, CU-02, CU-03, Green line, CU-04, Alavy green, CU-13, CU-14, and All Rounder. These genotypes generally showed lower fruiting intensity and yield, with some displaying less favorable phenological or postharvest responses. For example, CU-02 and CU-03 had relatively high fruit weight but weaker fruiting intensity, while Check variety Green line and Alavy green showed less favorable profiles for plant mortality or postharvest loss. The clustering further indicated that high yield was associated with a broader trait complex, particularly greater fruit number and harvest frequency, rather than with fruit size alone. This pattern was consistent with the correlation and PCA results and reinforced the favorable multivariate positions of CU-09, CU-10, and CU-11. From a breeding perspective, CU-09 and CU-10 are promising for prolific fruiting and high yield, whereas CU-11 offers a useful combination of productivity, heavier fruits, and thicker pericarp. Thus, the cluster analysis highlights both overall genotype similarity and complementary breeding value across the germplasm.

**Fig 3.**
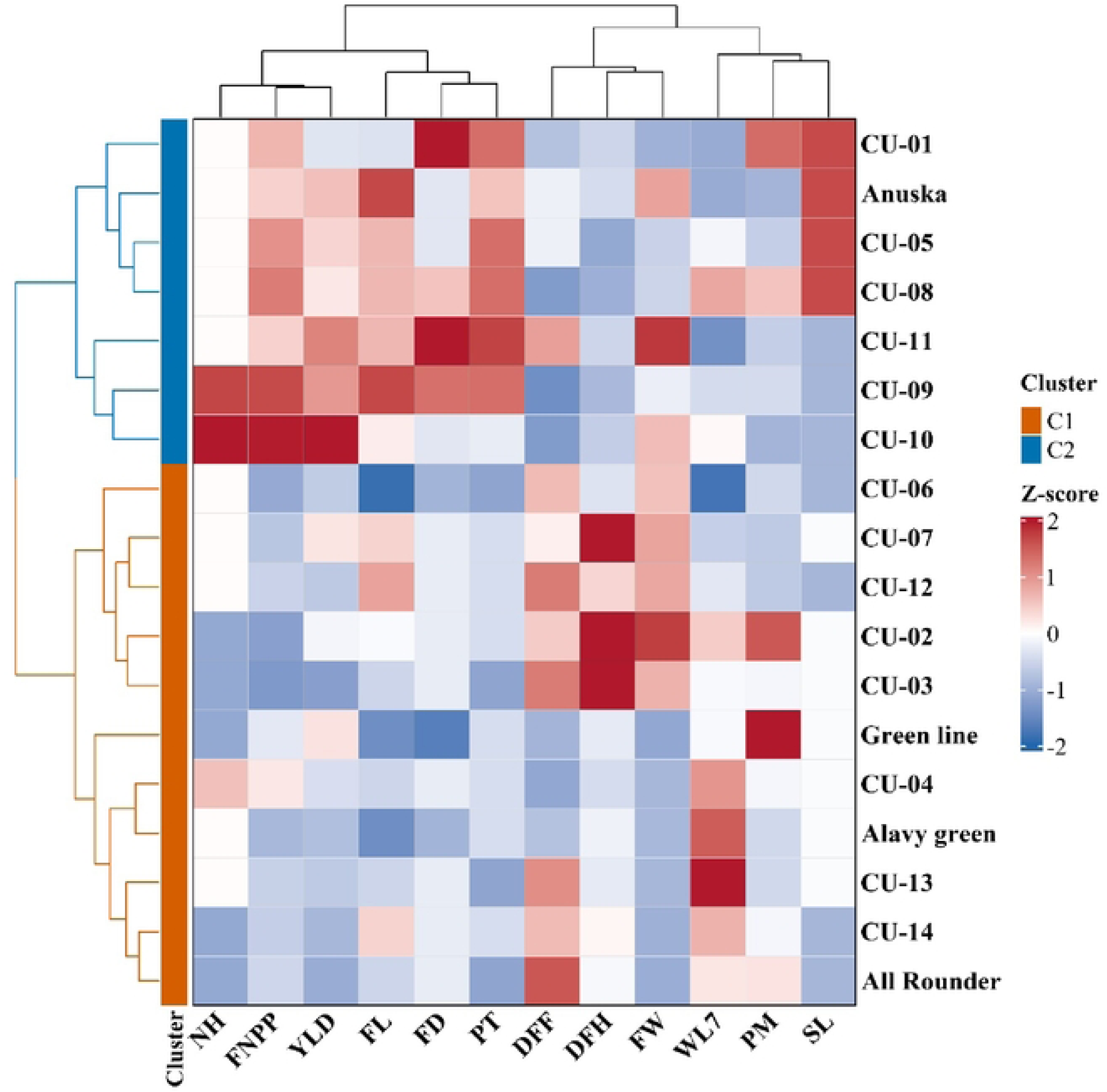
Hierarchical clustering heatmap illustrating the relationships among 18 cucumber genotypes based on 12 agronomic, yield, and postharvest traits. The heatmap was constructed using Z-score-standardized values of pooled adjusted genotype means across two years. Each row represents an individual cucumber genotype, while each column corresponds to a studied trait. The color gradient indicates relative trait values, with red representing higher values, blue indicating lower values, and white representing intermediate values. The dendrograms illustrate hierarchical relationships among genotypes and traits based on their multivariate profiles. The genotypes were classified into two major clusters: Cluster 1 (C1), comprising 11 genotypes and represented by orange annotations, and Cluster 2 (C2), comprising seven genotypes and represented by blue annotations. Differences in heatmap color patterns highlight variation in trait performance among the cucumber genotypes. **Note:** PM: Plant mortality; DFF: Days to female flowering; DFH: Days to first harvest; FL: Fruit length; FD: Fruit diameter; NH: Number of harvests; FNPP: Average fruit number per plant; FW: Average fruit weight; YLD: Yield; PT: Pericarp thickness; SL: Shelf life; WL7: Weight loss after 7 days; C1: Cluster 1; C2: Cluster 2.

### Multi-trait genotype selection based on performance and stability

Multi-trait selection further differentiated the cucumber genotypes by integrating performance across the 12 core traits (**Figure 4**). The MGIDI selection pattern strongly corroborated the single-trait evaluations while providing a more balanced multi-trait resolution. In individual trait assessments, CU-09 excelled in earliness and fruit length (15.00–16.70 cm), while CU-11 was distinguished by fruit diameter (5.00–5.60 cm) and single fruit weight (up to 136.20 g). When all 12 core traits were integrated into the genotype–ideotype distance framework, CU-09 and CU-11 (alongside the check variety Anuska) were selected by MGIDI due to their close proximity to the desired breeding ideotype (**Figure 4A**). Trait-wise factor loadings, communalities, factor assignments, and selection directions are provided in **Supplementary Table S3**. The complete MGIDI ranking and selection status of all 18 genotypes are presented in **Supplementary Table S4**. Interestingly, while CU-10 recorded the highest individual fruit yield (6.90–7.69 t ha⁻¹) and fruit number per plant (2.81–3.13) in univariate analyses, it was not retained by MGIDI, reflecting the index’s capacity to prioritize lines that achieve multi-trait equilibrium over extreme single-trait performers. In contrast, the MTSI analysis, which additionally accounted for relative interannual stability, selected CU-11, CU-01, and CU-06 (**Figure 4B**).

**Fig 4.**
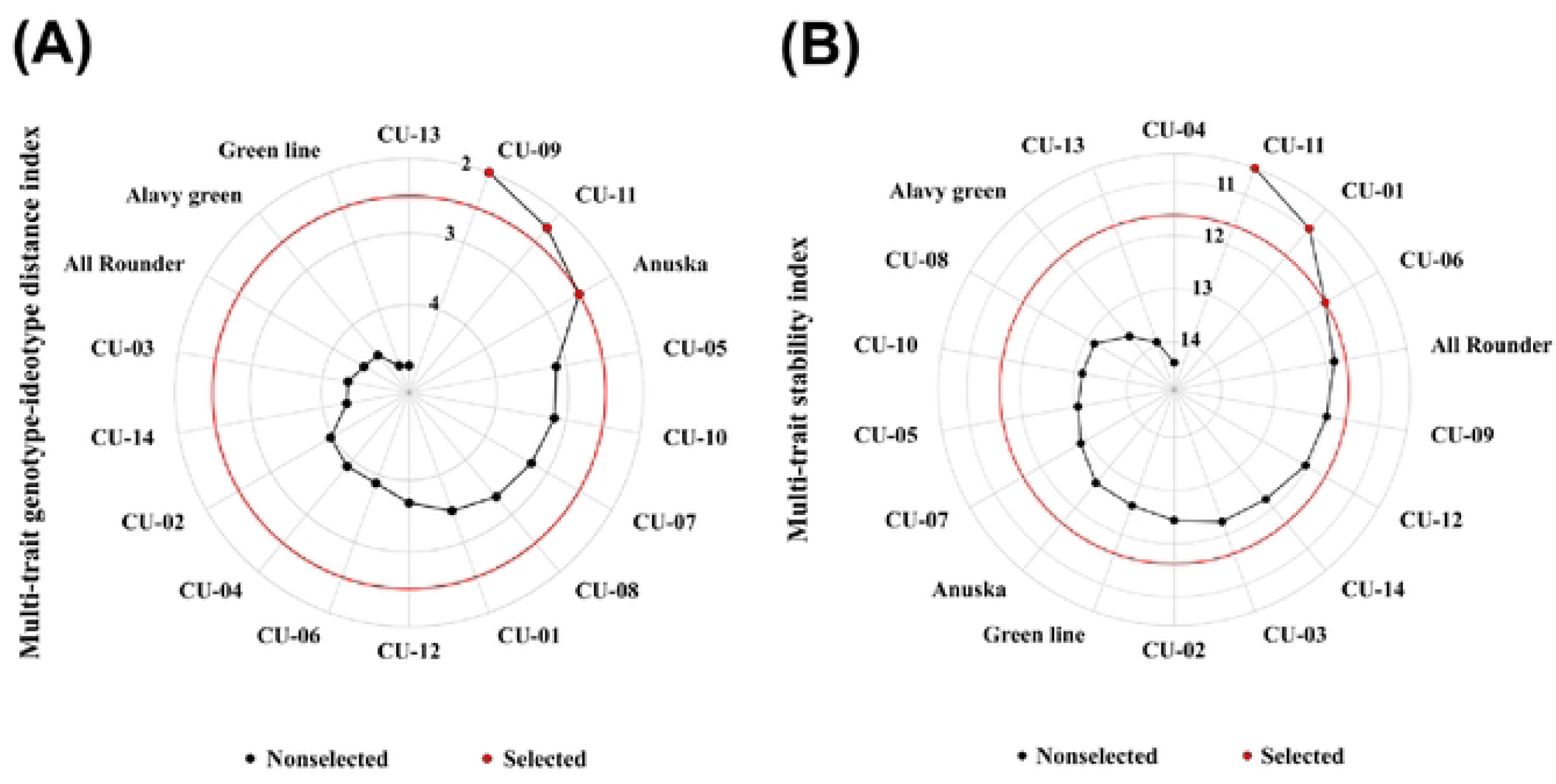
Multi-trait selection of cucumber genotypes based on overall performance and trait stability. (A) Multi-trait genotype–ideotype distance index (MGIDI)-based circular ranking of 18 cucumber genotypes using 12 core traits to identify genotypes with desirable multi-trait performance. (B) Multi-trait stability index (MTSI)-based circular ranking of the same genotypes to identify those combining desirable trait performance with stability. In both panels, red dots represent selected genotypes, whereas black dots indicate non-selected genotypes. The red circles indicate the selection thresholds based on the predefined selection intensity. Genotypes with lower index values are considered more desirable according to their respective selection criteria.

CU-11 was the only genotype selected by both indices, indicating a favourable combination of ideotype proximity and relative interannual stability within the two-year evaluation. The divergence between the remaining MGIDI and MTSI selections reflects the differing criteria of the two indices overall ideotype conformity versus temporal stability rather than contradictory performance. Taken together, MGIDI and MTSI provided complementary perspectives for identifying promising germplasm beyond yield alone. Representative fruits of six contrasting cucumber genotypes are presented in **Figure 5**, visually illustrating phenotypic variation in fruit length, diameter, and external appearance among check variety Green Line, Anuska, CU-05, CU-10, CU-09, and CU-11.

**Fig 5.**
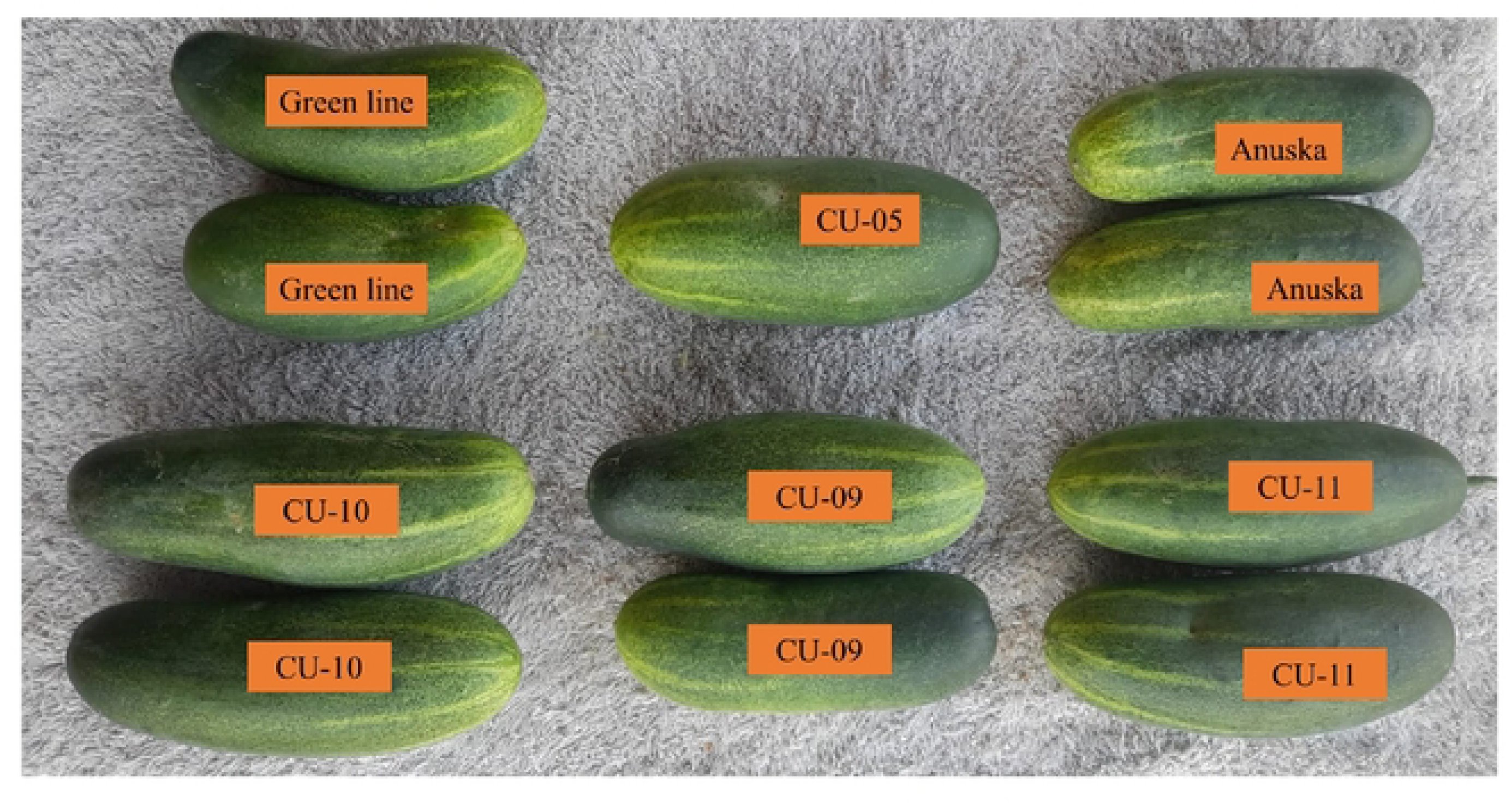
Representative fruit morphology of six contrasting cucumber (*Cucumis sativus* L.) genotypes evaluated during the two-year field experiment. The photograph illustrates visible variation in fruit size, shape, and external appearance among check variety Green line (upper left), CU-05 (upper center), check variety Anuska (upper right), CU-10 (lower left), CU-09 (lower center), and CU-11 (lower right). These representative fruits provide a visual comparison of morphological diversity among the evaluated cucumber germplasm.

## Discussion

### Genotypic Variation and Stability Across Years

Analysis of variance (ANOVA) showed highly significant genotype effects (P<0.001) for nearly all evaluated traits in both years, demonstrating substantial phenotypic differentiation among the 18 cucumber genotypes. The strong genotype effects for yield, fruit number plant^− 1^, and fruit weight indicate that between-genotype variation was much greater than residual variation for these economically important traits. Rani et al. [21] and Kumari et al. [22] likewise reported substantial genotypic variation for cucumber yield and associated traits, confirming substantial exploitable diversity within breeding materials. However, Singathiya et al. [23] demonstrated that genotype × environment interactions can alter trait expression across contrasting production environments, highlighting the importance of multi-environment validation. The consistency of significant genotype effects across both years further suggests that the ANOVA detected reproducible genetic differences rather than variation arising predominantly from experimental error.

### Genotypic mean performance for phenology and fruit traits

Marked and repeatable differences among the genotypes for mortality, flowering, harvest maturity, fruit dimensions, fruit number, and fruit weight indicate contrasting developmental and reproductive strategies within the evaluated cucumber germplasm. CU-10 and check variety Anuska maintained the lowest mortality (3.50–4.00%), whereas check variety Green line reached 24.00–26.80%, suggesting genotypic differences in establishment ability and early-season stress tolerance. Similar genotype-dependent variation in establishment has been reported in cucumber, although Singathiya et al. [23] showed that genotype × environment interaction can modify the magnitude of such responses under contrasting production conditions. Earliness also differed strongly, with CU-09 flowering in 16–18 days and CU-05 reaching first harvest in 34–38 days, whereas All-rounder, CU-02, CU-13, and CU-07 were considerably later. This agrees with Bisht et al. [24], who reported substantial variation and combining ability for earliness and yield-contributing traits in cucumber. Fruit morphology showed equally clear differentiation, with CU-09 and check variety Anuska producing the longest fruits and CU-01 and CU-11 the widest, reflecting genetic variation in longitudinal versus radial fruit growth. The most important yield-component contrast involved fruit number and fruit weight: CU-10 and CU-09 produced the most fruits plant^−1^, whereas CU-02 and CU-11 were characterized by heavier individual fruits. This pattern is consistent with source–sink competition, and recent cucumber studies similarly identified fruit number and fruit weight as important yield-associated traits [25]. Rather than indicating a rigid trade-off, the results show alternative yield architectures, because CU-09 combined high fruit set with long fruits, while CU-11 combined heavy fruits with greater diameter. Overall, CU-10 appears valuable for low mortality and prolific fruiting, whereas CU-09 combines earliness, elongated fruit shape, and high fruit set, making both useful parental materials for subsequent crossing and multi-environment validation.

### Yield and postharvest performance

Yield differed markedly among genotypes, with CU-10 consistently producing the highest yield (6.90–7.69 t ha⁻¹), followed by CU-11 and CU-09, whereas CU-03 remained the lowest-yielding genotype across both years. This advantage likely reflects differences in reproductive sink strength and assimilate allocation, consistent with Dhall et al. [5], who identified fruit number and fruit weight as major determinants of cucumber yield. However, Singathiya et al. [23] reported substantial genotype × environment effects for yield and quality traits, indicating that the consistently high yield of CU-10 across the two evaluated years warrants further multi-environment validation. Pericarp thickness also differed substantially, with CU-11 maintaining the thickest fruit wall (1.25–1.39 cm), which may improve mechanical strength and resistance to moisture loss during handling. Postharvest weight loss showed stronger genotypic discrimination than shelf life, with CU-06 and CU-11 recording the lowest losses, whereas CU-13 and check variety Alavy green exhibited the greatest dehydration, probably reflecting differences in cuticular resistance and epidermal permeability. In contrast, shelf life showed little differentiation in Year 1 but clearer separation in Year 2, suggesting stronger sensitivity to preharvest and seasonal conditions affecting respiration, water status, and senescence. The greater year dependence of shelf life is biologically plausible because cucumber storability is a complex trait influenced by fruit firmness, water loss, senescence, and other postharvest processes [26,27]. The contrasting profiles of CU-10 and CU-11 are particularly relevant because CU-10 combined maximum yield with moderate postharvest performance, whereas CU-11 combined high yield, heavy fruits, a thick pericarp, and relatively low weight loss. These results support simultaneous selection for productivity and postharvest durability, with CU-10 serving as a strong yield donor and CU-11 providing a more balanced source of yield and fruit-quality attributes for breeding.

### Genetic variability, heritability and selection potential

The narrow gaps between GCV and PCV for plant mortality, yield, fruit number plant^−1^, fruit weight, and 7-day weight loss indicate that environmental variation contributed relatively little to phenotypic expression of these traits. Their high broad-sense heritability further supports strong genetic control, although this parameter includes additive, dominance, and epistatic components and should not be interpreted as evidence of additive inheritance alone. The particularly high heritability and GAM for yield, fruit number, and fruit weight indicate strong scope for selection, consistent with recent cucumber studies reporting substantial genetic variability and selection potential for these productivity traits [25,28]. Bisht et al. [24] also reported an important contribution of non-additive gene action for yield-related traits, suggesting that hybrid breeding may complement direct phenotypic selection where dominance effects are substantial. Days to female flowering and first harvest also combined high heritability with high GAM, indicating that earliness can respond effectively to selection under the evaluated conditions. Plant mortality showed the greatest variability, heritability, and GAM, suggesting considerable opportunity to improve field establishment by selecting against genotypes with high stand loss. Fruit length and diameter also exhibited high heritability and high GAM, although their expected genetic gains were comparatively lower than those of yield and the major yield components. Weight loss after 7 days also showed high heritability and appreciable GAM, supporting the feasibility of improving postharvest moisture retention through selection. In contrast, shelf life exhibited low GCV, moderate heritability, low GAM, and genotype × year variance exceeding genotypic variance, indicating strong environmental dependence and reduced reliability of single-season selection. Collectively, yield, fruit number, fruit weight, earliness, plant survival, and postharvest weight retention are comparatively strong selection targets, whereas shelf life, pericarp thickness, and fruit dimensions require broader multi-season or multi-environment validation before stable breeding decisions are made.

### Interannual Stability and Trait-Specific Environmental Sensitivity

Fruit number plant^−1^, yield, and plant mortality showed the strongest interannual repeatability, indicating highly stable genotype rankings for reproductive capacity and field survival across years. This stability is biologically consistent with strong genetic control over fruit set, assimilate partitioning, and establishment-related traits, which reduces the likelihood of major rank changes under moderate seasonal variation. Comparable temporal and environmental variability in cucumber yield and earliness has been reported in multi-year evaluations [6,29], while fruit structural traits such as length, diameter, and pericarp thickness generally show greater environmental responsiveness. The lower repeatability of these structural traits, and especially the non-significant shelf-life correlation (r=0.426; P=0.078), likely reflects stronger sensitivity to preharvest temperature, moisture status, and postharvest senescence processes, which can override genotypic differences between years. These findings suggest relatively consistent genotype rankings for several productivity traits between the two study years, whereas fruit morphology and particularly shelf life require further validation across years and locations.

### Trait Associations and Breeding Implications

Yield was more strongly associated with fruit number plant^−1^ and harvest frequency than with fruit weight, indicating that cumulative fruiting capacity was more strongly associated with productivity than individual fruit weight. Biologically, a larger number of active reproductive sinks sustains assimilate demand across successive harvests, which agrees with Patel et al. [2] and Subedi et al. [6], who identified fruit number as a major determinant of cucumber yield. Earlier flowering and harvesting were also associated with higher fruit number and yield, while their positive association with fruit weight suggests that later genotypes may partially compensate for fewer fruits through greater assimilate allocation to individual fruits. This earliness–fruit-size relationship supports selection for rapid reproductive transition when cumulative yield is prioritized over maximum individual fruit size. Fruit length, diameter, and pericarp thickness became more strongly associated with yield in the second year, indicating greater environmental responsiveness of structural fruit traits than of core fruiting traits. Such seasonal shifts likely reflect the sensitivity of cell expansion and pericarp development to assimilate availability and plant water status, which may explain differences from studies reporting more stable morphological associations [12,13]. Postharvest weight loss remained negatively associated with fruit weight, whereas shelf-life relationships changed markedly between years, supporting stronger environmental control of storability [8]. The negative associations of plant mortality with fruit number, harvest frequency, fruit weight, and yield further indicate that strong field establishment supports reproductive performance. Overall, simultaneous selection for earliness, prolific fruiting, repeated harvest capacity, and low mortality is likely to be more effective than selection for fruit size alone, while shelf life requires multi-season validation.

### Multivariate analysis of genotype differentiation and hierarchical structure

Principal component analysis and hierarchical clustering consistently separated the cucumber genotypes according to coordinated differences in earliness, fruiting intensity, yield, fruit morphology, and pericarp characteristics. The main multivariate gradient contrasted genotypes with high fruit number, repeated harvesting, and greater yield against those with later flowering and harvest maturity, reflecting differences in reproductive sink establishment and assimilate allocation. Hierarchical clustering reinforced this structure by separating the 18 genotypes into two major groups, with Cluster 2 containing CU-01, check variety Anuska, CU-05, CU-08, CU-11, CU-09, and CU-10, which generally expressed higher fruit number, harvest frequency, yield, and more favourable quality attributes, whereas Cluster 1 contained genotypes with weaker fruiting intensity or less favourable phenological and postharvest profiles. Comparable multivariate grouping has been reported by Kumar et al. [13], Ene et al. [1], Patel et al. [2], and Acharya et al. [3], who likewise identified yield components and phenological traits as major contributors to cucumber diversity. The agreement among these studies likely arises because flowering time, fruit set, fruit growth, and yield are physiologically interconnected and therefore generate coordinated covariance patterns that are captured by both ordination and clustering methods. Within the higher-performing cluster, CU-09 and CU-10 were characterized mainly by prolific fruiting and high yield, whereas CU-11 combined productivity with heavier fruits and thicker pericarp, demonstrating that superior performance can arise through different but complementary trait architectures. The broad correspondence between principal component ordination and hierarchical clustering therefore strengthens the evidence that genotype divergence was driven by integrated trait complexes rather than isolated characters, supporting their combined use for identifying complementary parents and reducing redundancy in breeding populations.

### Multi-trait genotype selection based on performance and stability

A major challenge in conventional vegetable breeding is that selection based purely on single-trait means can result in undesirable correlated responses due to trait trade-offs. Comparing univariate genotypic means with the multi-trait genotype–ideotype distance index (MGIDI) in this study highlights this dynamic. The selection of CU-09 and CU-11 by MGIDI confirms that their superior univariate attributes namely early female flowering, elongated fruit morphology, and prolific fruiting in CU-09, and heavy, thick-walled fruits with low postharvest weight loss in CU-11 collectively conform to the ideal cucumber ideotype (Fig 4A). In contrast, genotypes that ranked foremost in single yield parameters, such as CU-10 (highest yield and fruit number per plant) and CU-05 / CU-08 (earliest harvest maturity), were not selected under the multi-trait index. In the case of CU-10, exceptionally high reproductive sink demand coincided with only moderate postharvest durability and smaller fruit dimensions, which increased its multivariate distance from the ideal ideotype. Similarly, the early harvest readiness of CU-05 and CU-08 did not sufficiently offset lower ratings in fruit weight or postharvest stability. These discrepancies demonstrate that MGIDI effectively prevents the over-selection of single-trait extremes, confirming CU-11 and CU-09 as the most balanced, ideotype-proximate advanced lines for cucumber population improvement and parent selection. Furthermore, the divergence between MGIDI and MTSI highlights that proximity to a desirable multi-trait ideotype does not necessarily imply consistent trait expression across years. CU-11 was the only genotype retained under both selection approaches, demonstrating a unique combination of ideotype proximity and interannual stability. In contrast, CU-09 and check variety Anuska were favored under the ideotype-based performance criterion, whereas CU-01 and CU-06 were preferred when relative interannual stability was additionally considered. These differences reflect the complementary objectives of ideotype-based and stability-oriented multi-trait selection [30,31]. Nevertheless, the two-year evaluation at a single location provides only preliminary evidence of stability, emphasizing the need for additional multi-environment trials before recommending genotypes for broader adaptation.

## Conclusions

The two-year evaluation revealed substantial and repeatable genetic variation among 18 cucumber genotypes for agronomic, yield, fruit-quality, and postharvest traits. Yield, fruit number plant^−1^, and fruit weight showed high genetic variability, heritability, and genetic advance, confirming their value as primary selection criteria. CU-10 was consistently the highest-yielding genotype, whereas CU-11 combined high yield with heavier fruits, thicker pericarp, and lower postharvest weight loss. Strong interannual repeatability for major productivity traits further indicated that genotype rankings were largely stable across the two evaluation years. Multi-trait analyses identified CU-11 as the only genotype selected by both MGIDI and MTSI, indicating a favorable combination of multi-trait performance and relative interannual stability. PCA and hierarchical clustering also consistently differentiated superior genotypes on the basis of coordinated variation in earliness, fruiting intensity, yield, and fruit-quality traits. Overall, CU-11 is a promising candidate for further evaluation and parental use due to its balanced performance and relative interannual stability. CU-09 may contribute favorable earliness and fruit-shape traits, while CU-10 represents a potential source of high yield and harvest frequency. Anuska, CU-01, and CU-06 may provide complementary breeding value. The integrated use of quantitative genetic parameters and multivariate selection therefore offers a robust framework for identifying superior cucumber germplasm. Further multi-location and multi-season validation is required before broad cultivar recommendation.

## Limitations

This study was conducted at a single location over two consecutive Rabi seasons, which limits inference regarding genotype performance across broader agro-ecological environments. Several traits, particularly shelf life and pericarp-related attributes, showed greater year-dependent variation, indicating the need for validation across additional seasons and locations. Moreover, the study focused primarily on phenotypic and quantitative genetic parameters; integrating molecular marker data and controlled postharvest assessments would strengthen future genotype characterization and selection.

## Acknowledgments

Not applicable

## Author contributions

Conceptualization: Mohaimin Islam.

Data curation: Md. Omar Kayess, Md Salman Shahriar.

Formal analysis: Md. Omar Kayess, Md Salman Shahriar.

Investigation: Mohaimin Islam, Md. Omar Kayess, Md Salman Shahriar

Methodology: Md. Omar Kayess

Project administration: Md. Omar Kayess

Software: Md. Omar Kayess, Md Salman Shahriar. Supervision: Md. Omar Kayes

Validation: Md. Omar Kayess.

Visualization: Md. Omar Kayess.

Writing – original draft: Md. Omar Kayess, Md Salman Shahriar, Mst. Faria Kabir, Md. Anwar Hossain.

Writing – review & editing: Md. Omar Kayess, Md Salman Shahriar.

## Ethics statement

Not applicable

## Supplementary Files Captions

Supplementary Table 1. Mean Monthly meteorological conditions during the Rabi-season cucumber growing period in the Bogura region, Bangladesh in both years

Supplementary Table 2: List of Experimental Genotypes and Check Varieties Evaluated in the Study

Supplementary Table 3. Trait-wise factor assignments, observed means, selection means, and selection differentials for 24 traits used in the multi-trait genotype–ideotype distance index (MGIDI) analysis.

Supplementary Table 4. Ranking and Selection of 18 Genotypes Based on the Multi-Trait Genotype– Ideotype Distance Index (MGIDI).

